# The giant pouched rat (*Cricetomys ansorgei*) olfactory receptor repertoire

**DOI:** 10.1101/742668

**Authors:** Angela R. Freeman, Alexander G. Ophir, Michael J. Sheehan

## Abstract

For rodents, olfaction is essential for locating food, recognizing mates and competitors, avoiding predators, and navigating their environment. It is thought that rodents may have expanded olfactory receptor repertoires in order to specialize in olfactory behavior. Despite being the largest clade of mammals and depending on olfaction relatively little work has documented olfactory repertoires outside of the conventional laboratory mice and rats. Here we report the olfactory receptor repertoire of the African giant pouched rat (*Cricetomys ansorgei*), a Muroid rodent distantly related to mice and rats. The African giant pouched rat is notable for its large cortex and olfactory bulbs relative to its body size compared to other sympatric rodents, which suggests anatomical elaboration of olfactory capabilities. We hypothesized that in addition to anatomical elaboration for olfaction, these pouched rats might also have an expanded olfactory receptor repertoire to enable their olfactory behavior. We examined the composition of the olfactory receptor repertoire to better understand how their sensory capabilities have evolved. We identified 1145 functional olfactory genes, and 260 additional pseudogenes within 301 subfamilies from the African giant pouched rat genome. This repertoire is similar to mice and rats in terms of size, pseudogene percentage and number of subfamilies. Analyses of olfactory receptor gene trees revealed that the pouched rat has 6 expansions in different subfamilies compared to mice, rats and squirrels. We identified 99 orthologous genes conserved among 4 rodent species and an additional 167 conserved genes within the Muroid rodents. The orthologous genes shared within Muroidea suggests that there may be a conserved Muroid-specific olfactory receptor repertoire. We also note that the description of this repertoire can serve as a complement to other studies of rodent olfaction, as the pouched rat is an outgroup within Muroidea. Thus, our data suggest that African giant pouched rats are capable of both natural and trained olfactory behaviors with a typical Muriod olfactory receptor repertoire.

## Introduction

In rodents, olfaction is essential for a number of behaviors including social recognition (1), sexual behavior (2), predator detection (3), and finding food (4). The mechanism for olfactory perception in rodents uses two systems, the main olfactory system and the vomeronasal system, the former of which contains the olfactory receptors (ORs) which transduce odorants into a perceived ‘smell’ (5). The repertoire of these ORs has been of particular interest, as the OR gene family is one of the largest in the genome, and the overall number of functional receptor genes is thought to correlate with olfactory perceptual ability (6). In particular, expansion and contraction of the OR repertoire (i.e. functional OR genes) among species is thought to correlate with changing environmental niche – e.g., animals that use vision primarily for navigation and recognition tend to have more pseudogenes compared to functional ORs (7–9). Furthermore, certain families of ORs are associated with diet and niche, with larger repertoires generally associated with asocial, terrestrial, and diurnal species [(10,11) but see (12)].

ORs are G-protein coupled receptors that are located in olfactory sensory neurons (OSN) of the nasal epithelium, where they bind odorants to transmit information to the brain (13). Which ORs bind which odorants is still largely unknown, as there are a number of technical limitations for high-throughput screening methods for ligand identification (14,15). Despite these limitations, research shows that related ORs within subfamily groups (and orthologous OR genes among taxa) tend to bind structurally similar ligands (16), though the selectiveness and level of response by individual OSNs may vary (9,17–21). When an odorant binds to the OR in the OSN, a signal travels down that neuron’s axon. The axon projects onto glomeruli in the olfactory bulb, which allows for the signal to be transmitted and interpreted. Although OSNs only contain one type of OR, the OR can bind a range of odorants causing this olfactory information to be coded using a combinatorial method (21). For example, a molecule might activate OSNs only in specific changing concentrations, or in mixtures (or alone) (5,22). This elegant system of many concurrently activated or inactivated ORs in combination is what allows animals to perceive their olfactory-world and respond accordingly.

To understand how evolution may have influenced both the number and distribution of ORs, several studies have been conducted on the OR repertoire across species. Since the first description of ORs in mice by Axel and Buck (1991) (13), OR genes in mammals have been more extensively described in mice (*Mus musculus*) (23), but also in swine (*Sus scrofa*) (24), humans (*Homo sapiens*) (17), dogs (*Canis familiarisi*) (25,26), cattle (*Bos taurus*) (27), non-human primates (8,12), guinea pig (*Cavia porcellus*), rabbit (*Oryctolagus cuniculus*), horse (*Equus caballus*), and elephant (*Loxodonta africana*) (28), among other mammals (11).

By investigating the overall diversity of OR gene repertoires, we can further elucidate how this large receptor family has evolved and whether these repertoires have adapted over time for certain environments and behavior specializations. Certainly, rodents are expected to have large OR repertoires, because much of their behavior relies on olfaction. However, OR repertoires of rodents still vary; guinea pigs are reported to have fewer than 1000 OR genes (functional and pseudogenes), while other mammalian species have olfactory repertoires twice that size (11). In order to expand our understanding of olfaction and evolution within mammals, we have examined the African giant pouched rat (*Cricetomys ansorgei*) olfactory genome.

The African giant pouched rat (colloquially, ‘Pouchie’) is a large, mostly nocturnal rodent native to the savannah of Africa (29). *Cricetomys* are within Nesomyidae which shared a common ancestor with Muridae rodents (i.e. *Mus* and *Rattus*) approximately 18-20 MYA (**Fig 1**) (30,31). The name ‘pouched rat’ is a misnomer given that it is only a distant relative of the conventional *Rattus*. Furthermore, the term ‘African giant pouched rat’ can refer to any of the 6 species of *Cricetomys* (*C. ansorgei, C. emini, C. gambianus, C. kivuensis*, and two unnamed species) (32). The *Cricetomys* genus is widely distributed across sub-Saharan Africa, and different species tend to associate in different environmental niches (32). A uniting feature of this genus and other members of the subfamily Cricetomyinae (i.e. *Beamys* and *Saccostomus*) (33) are the gerbil-like cheek pouches where *Cricetomys* store food during foraging. However, their physical resemblance to *Rattus* is most likely a result of convergent evolution from distinct ancestors. *Cricetomys* species have notably large olfactory bulbs and neocortex for their size - even when compared to other rodents (29,34). The olfactory bulbs of the pouched rat comprise 19% of the total brain length, while the greater cane rat (*Thryonomys swinderianus*) have olfactory bulbs of only 9% of the brain, despite occupying similar regions in Africa (34,35). Furthermore, the neocortex for the pouched rat accounts for 75% of the cerebral cortex (compared to 50% in mice) (35), and the neocortex ratio places the pouched rat in the range of primates (29). The anatomical features of the olfactory system can predict the size of the OR repertoire (36,37), thus, we predicted that the pouched rat may also have an expanded OR repertoire.

**Fig 1.**
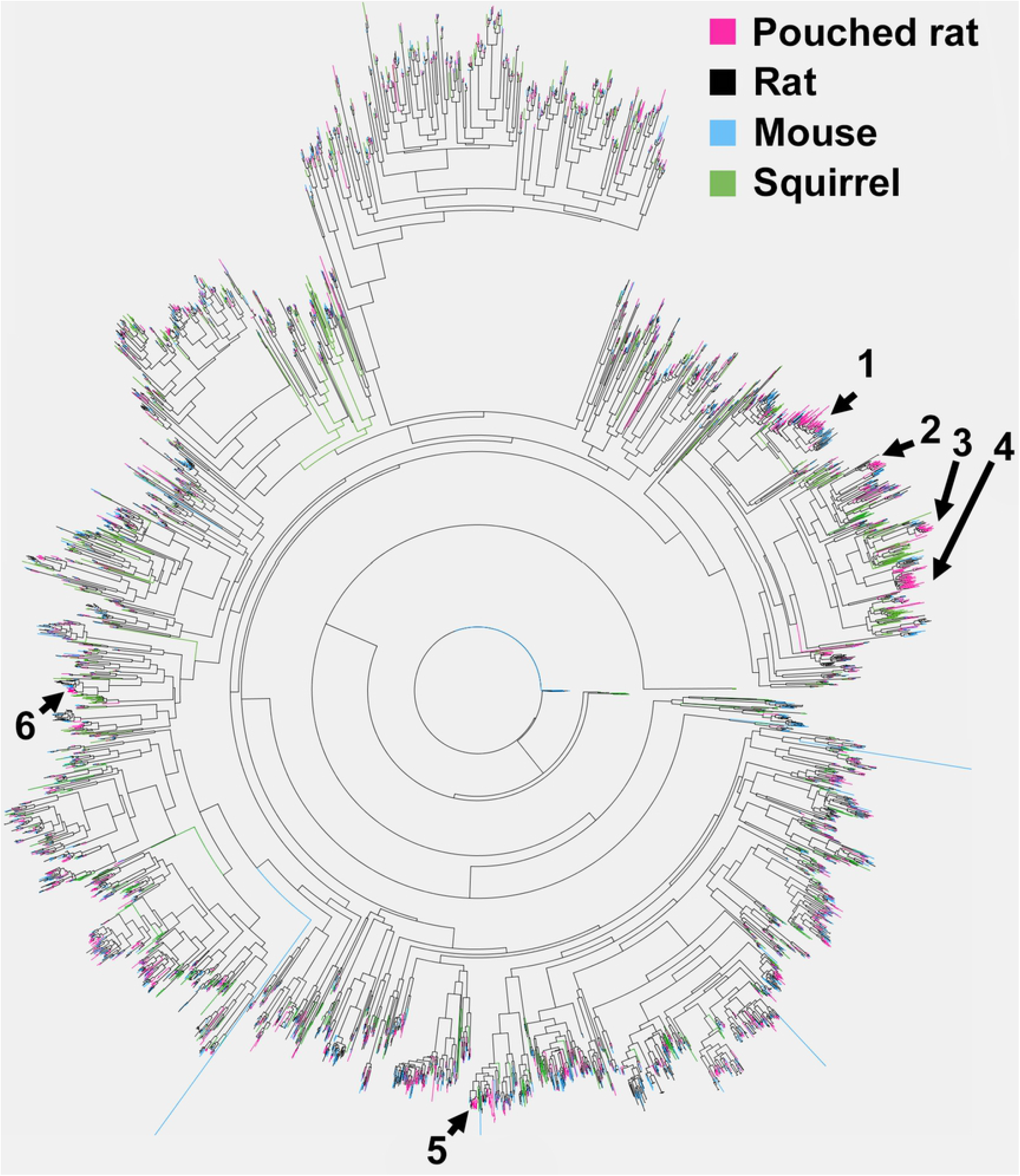
Phylogeny of species compared in this study. The figure shows a simplified phylogeny of mice (*Mus sp*.), rats (*Rattus sp*.), pouched rats (*Cricetomys sp*.), and squirrels (*Ictidomys sp*.). Pouched rats are Nesomyid rodents that are distantly related to Murids. Silhouettes show approximate size differences among species. Time periods are listed along the x-axis. MYA: million years ago, Plioc: Pliocene, IV: Quaternary period.

In addition to these anatomical features, recent behavioral research emphasizes the pouched rats’ olfactory ability. Pouched rat males can use olfactory cues to discriminate between reproductively available and unavailable females, and similarly, females exhibit preferences for ‘competitive’ males based on olfactory cues (38). Furthermore, males can perceive olfactory cues from other males’ scent marks which they countermark in unfamiliar contexts (39). Pouched rats are even used as ‘biodetectors’ and can be trained to detect tuberculosis and TNT via olfaction (40,41). Given their anatomical adaptations for olfaction, their use of olfactory cues in potential competitor and mate assessment, and their use as olfactory biodetectors, we assessed the olfactory repertoire of *C. ansorgei* to determine how the pouched rat OR repertoire compares to other rodents. We hypothesized that in addition to anatomical elaboration of the olfactory system, pouched rats would have an expanded OR repertoire which enables their olfactory behavior.

Most rodents that have been previously studied for their OR repertoire, or used for comparative work, are limited to lab models (i.e. Guinea pig, rat (*Rattus norvegicus*), mouse) (26,28), though recently some others have been described (11). Thus, our aim was to describe the olfactory receptor repertoire of *C. ansorgei* as a representative of *Cricetomys*, which would expand the diversity within described *Rodentia* olfactory receptor gene repertoires. Here we describe ORs in this species and compare their distribution and number to other rodents (**Fig 1**).

## Methods

We downloaded the publicly available draft assembly of the pouched rat genome (NCBI ID 75238 (CriGam_v1_BIUU). This genome contains 1,102,952 scaffolds, and 1,110,425 contigs. The scaffold N50 and L50 values are 110,049 and 5,340, while the contig N50 and L50 are 80,761 and 7,273, respectively. The genome is reported to have 36.7x coverage and a GC% of 42.1%.

### Detection of OR genes in the pouched rat genome

We created a database of ORs from human (*H. sapiens*), mouse (*M. musculus*), rat (*R. norvegicus*), and squirrel (*Ictidomys tridecemlineatus*) from Ensembl BioMart (42), using the search term ‘olfactory receptor’ and the most recent genome versions. We then performed a translated basic local alignment search (TBLASTN) to identify potential ORs within the pouched rat genome. We included all matches, which had e-values of 0, and/or the match started at 1, protein alignment lengths of > 133 amino acids, bit scores of > 224, and gaps of 40 or fewer amino acids. These parameters set a threshold; below these parameters matches declined quickly in similarity. For mouse matches, this reduced the number of total results (1,148,779) to 1213, and for rat matches reduced the number of total results (1,219,637) to 1219. For human and squirrel matches, all the matches were above the threshold and included 903 and 35 sequences, respectively. We used the Galaxy web platform to conduct a multiple alignment using fast fourier transform (MAFFT) to locate and remove duplicate sequences and combine overlapping sequences (43). This resulted in 2,399 sequences which coded for potential pouched rat ORs. We then extended the ends of sequences by 100 bp to ensure that we would obtain the full coding region, and excluded any genes of less than 750 bp, because most functional OR genes are >250 amino acids (44), which resulted in 1409 sequences. We obtained all the open reading frames (via EMBOSS) (45) within all the sequences (~23k; 84% were shorter than 50 amino acids in length) and filtered out those shorter than 250 amino acids (44). This left 1,149 functional coding genes. Using these putative functional ORs, we generated a multiple sequence alignment (via MAFFT) and produced a phylogenetic tree using FastTree to examine any outliers. FastTree was chosen at this step instead of RAxML for processing efficiency (46). We used BLAST on any outliers (long branches separate from other groups) in the tree, to verify they encoded olfactory receptors, which led to the removal of 4 sequences which were most similar in terms of sequence similarity to other known G-coupled receptors. Thus, out of the 1405 potentially functional OR receptors, 18.5% were labeled ‘pseudogenes’ that likely did not produce a functional receptor product (but see (47)), either due to truncation, deletions or frameshift mutations. Using these ‘manual similarity-pipeline’ search and selection methods, we found 1145 putative functional OR receptor genes.

In addition to the above pipeline, we also used the Olfactory Receptor Assigner (ORA) module to detect potential OR genes and pseudogenes in the entire pouched rat genome (11). This method identified a similar number of functional OR genes, detailed methods are available in the supplement (S1 File). The ORA module detected 1060 unique putative functional OR receptor genes from the pouched rat genome.

### Classification based on phylogeny

We produced a phylogenetic tree using the 1145 putative functional pouched rat OR sequences located during the manual pipeline search using MAFFT (L-INS-I method) for sequence alignment and RAxML to produce an unrooted tree with 1000 bootstraps using the PROTCATJTTF option. We then used the method described by Zhang and Firestein (2002), to determine family categorization, by > 50% bootstrap support and 40% protein identity. Subfamilies were defined as >50% bootstrap support and 60% protein identity (48). Protein identity was determined through a BLASTP pairwise-comparison. To determine clades, high bootstrap support is typically used (44), the pouched rat olfactory repertoire separated into two clades with 76 and 71% bootstrap support.

### Comparative rodent OR phylogenetic tree

We used a MAFFT alignment (L-INS-I method) and RaxML to produce an unrooted tree with 500 bootstraps using the PROTCATJTTF option to compare the 1145 OR protein sequences of mouse, rat, squirrel, and pouched rat. This comparative rodent OR phylogenetic tree was used for examination of expansions and orthology.

### Expansions within gene families

We used the rodent OR phylogenetic tree to identify pouched rat-specific expansions. We compared pouched rat functional gene distributions (number of genes in the subfamily) and compared this to the most similar mouse, rat, and squirrel genes. For each noted expansion, we located the most closely related mouse OR gene and identified its subfamily group to determine if there were potential identified ligands within the mouse OR subfamily.

### Orthologous ORs

Using the rodent tree created from protein sequences, we examined the branching to identify orthologous ORs among pouched rats and squirrel, mouse, and rat. We enumerated instances where two paralogs (from any of the rodent species) were indicated in the tree. For larger numbers of paralogs (i.e. larger expansions), we did not include these in paralog or ortholog enumeration. The structure of branching for each ortholog group was recorded and compared to shared sequence identity. We then calculated the shared sequence identity for each ortholog compared to the pouched rat for each group.

### Clustering within the genome

We analyzed the relative positions of the putative functional pouched rat ORs obtained from the manual similarity-pipeline, by grouping gene clusters according to positional proximity. The pouched rat genome was parsed into a custom BLAST database (using makeblastdb (49)), which allowed for a conservative estimate of physical clustering – as sequences within the same parsed cluster of the genome would be physically clustered in a chromosome. OR genes were considered ‘clustered’ when they were located within the same sequence identifier.

## Results

We identified 1145 functional pouched rat ORs from the pouched rat genome using our manual pipeline method, which is similarly-sized compared to repertoires of mouse and rat (S2-S3 files). The manual pipeline method and ORA method described similar numbers of ORs, and details of the ORA method and results are available in the supplement (S1 File). We identified 260 pseudogenes with the manual pipeline method.

The pouched rat has similar numbers of clades, pseudogenes, or truncated genes to mice and rats, and has a relatively similar number of subfamilies to most other species using the manual pipeline search method (Table 1, **Fig 2, S4 file**).

**Table 1.** Distribution of pouched rat olfactory receptor genes among subfamilies using a manual pipeline search method. (48)

| Number of sequences in<br>subfamily | Number of instances |
| --- | --- |
| 1 | 140 |
| 2 | 48 |
| 3 | 25 |
| 4 | 24 |
| 5 | 10 |
| 6 | 11 |
| 7 | 4 |
| 8 | 6 |
| 9 | 5 |
| 10 | 5 |
| 11 | 3 |
| 13 | 1 |
| 14 | 2 |
| 15 | 5 |
| 16 | 2 |
| 18 | 2 |
| 19 | 1 |
| 22 | 1 |
| 24 | 1 |
| 26 | 1 |
| 31 | 2 |
| 32 | 1 |
| 49 | 1 |

**Fig 2.**
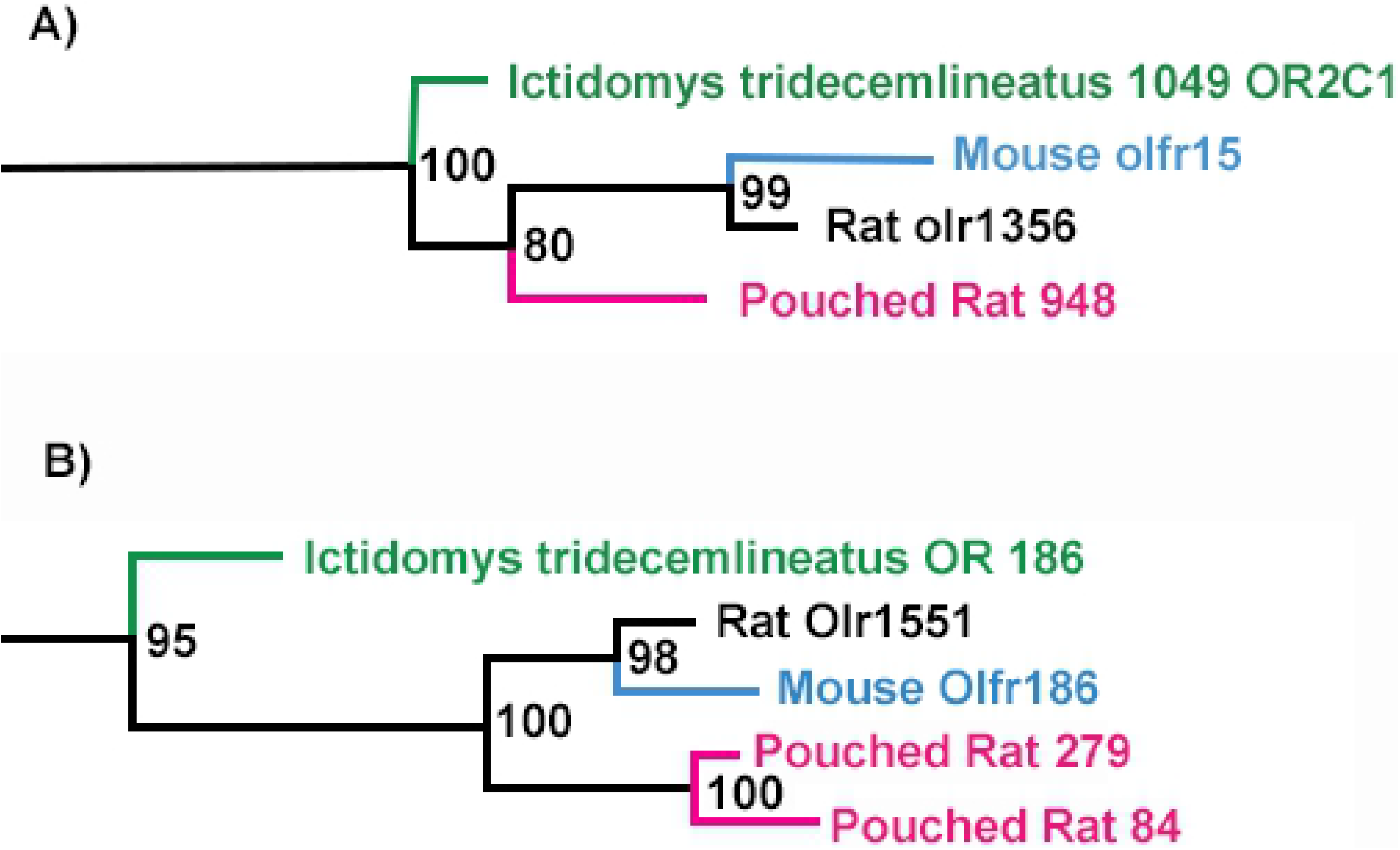
Family-group representation in the pouched rat olfactory receptor gene tree. Different colors at nodes indicate different families. All families share > 50% bootstrap support and 40% protein identity.

The pouched rat has similar numbers of functional and pseudogenes (**Fig 3**) compared to other rodents, and similar numbers of subfamilies to mice (Table 2). Ranges within table 3 indicate variation in reported values due to variation in search strategies.

**Fig 3.**
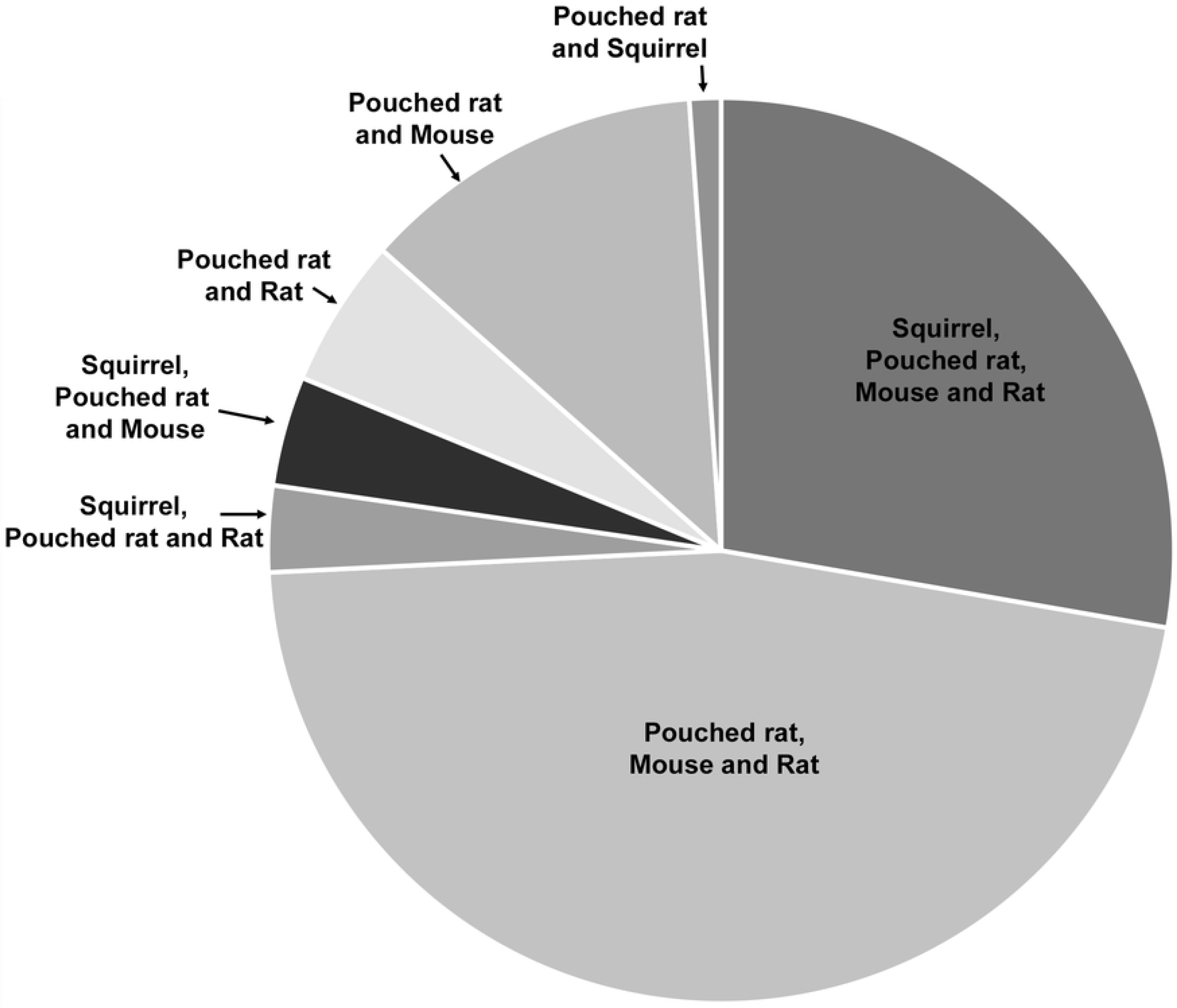
Graph comparing functional and pseudogene numbers among select species. Black bars indicate functional genes and grey bars indicate pseudogenes. Pouched rats, rats, mice,squirrels, dogs, cattle, and pigs have similar numbers of reported olfactory receptor genes. Squirrels have large numbers of pseudogenes.

**Table 2.** Families, subfamilies and functional olfactory receptor genes within selected species.

| Species | Number<br>of families | Number of<br>subfamilies | Number of<br>functional genes<br>(%) | Number of<br>pseudogenes |
| --- | --- | --- | --- | --- |
| African giant<br>pouched rat<br>(*C. ansorgei*), this<br>paper | 87 | 301 | 1145 (82) | 260 |
| Squirrel (11) | 16 | nd | 830 (32) | 1763 |
| Rat (6,26,27) | 21 | 282 | 1201-1217 (71-80) | 292 |
| Mouse (23,27,48) | 228 | 241 | 913-1037 (75-80) | 354 |
| Human (25,27,50,51) | 17-61 | 285-300 | 388-390 (48) | 414-465 |
| Dog (6,25-27) | 23 | 300 | 811-875 (75-82) | 222-278 |
| Pig (27) | 17 | 349 | 1113 (86) | 188 |
| Cattle (27) | 18 | 272 | 881 (82) | 190 |
| Zebrafish (27,52) | 8 | 40 | 102-143 (74-86) | 17-35 |
| Chicken (53,54) | nd | nd | 82-368 (15-66) | 111-476 |
| Frog (27) | nd | nd | 410 (46) | 478 |

nd: no data available

To compare pouched rat subfamilies with other species, we produced a gene tree using the sequences identified during manual pipeline search (**Fig 4**; S5 File), which illustrates that for most mouse, rat, and squirrel ORs subfamilies, there is at least one pouched rat OR (A tree using sequences from the ORA was visually similar in structure, and is provided in the supplement: S6 File). Of particular interest is the number and distribution of lineage specific gene family expansions. We can identify potential expansions where there are relatively more pouched rat ORs and fewer ORs from the other rodents within a family or subfamily. We have indicated these expansions, and their associated family groups in **Fig 4** for reference. The first expansion includes ORs from pouched rat subfamily 329 (S7 File); the largest pouched rat subfamily. It is associated with mouse OR subfamily 150 (Human equivalent Family 1, Subfamily E) (23). There are 6 ORs in the associated mouse subfamily, but 35 pouched rat ORs (23) (Table 4). The second expansion includes ORs from the pouched rat subfamily 312. This expansion was associated with mouse subfamily 137. The third and fourth expansions include ORs from pouched rat subfamily 321 and 317, respectively. Pouched rat subfamily 317 is the second largest pouched rat subfamily. The third expansion is associated with mouse subfamily 135 (Human equivalent Family 7 Subfamily E) and the fourth expansion is associated with mouse subfamily 136 (Human equivalent Family 7 Subfamily E). A fifth expansion includes pouched rat subfamily 12, which is related closely to mouse olfr183 and mouse subfamily 89, and also rat olr1555. The last noted expansion is associated with mouse subfamily 71. The distribution of ORs by subfamily size, as defined by (48) is shown in Table 2 and the composition of expansions are shown in Table 4 (comparative results from the ORA are shown in the Table in S1 Table).

**Fig 4.**
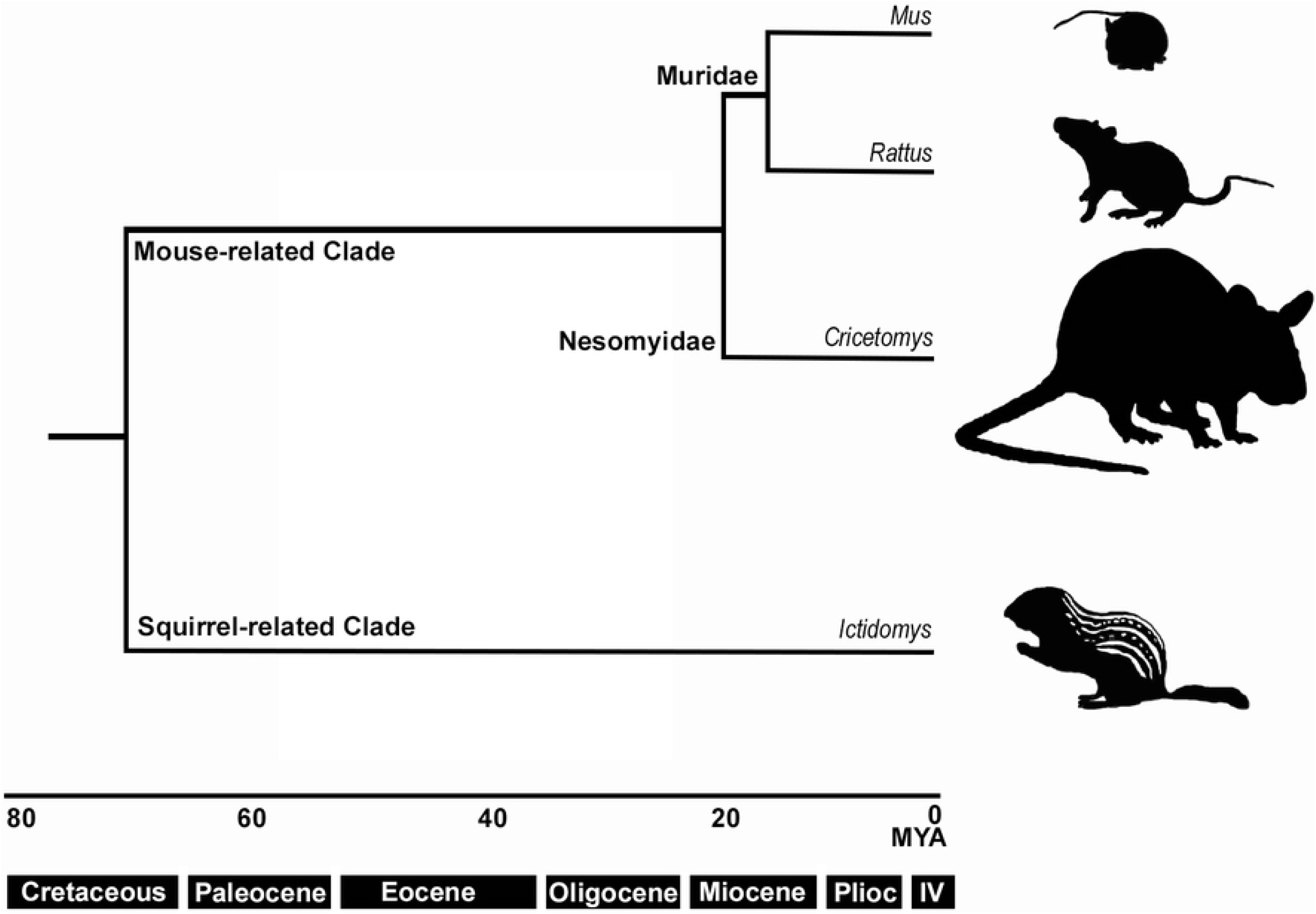
Comparative unrooted gene tree of olfactory receptor sequences for pouched rat, mouse, rat, and squirrel. Expansions are indicated with group identifiers and arrows.

**Table 4.** Expanded subfamilies of pouched rat olfactory receptors (ORs) and associated mouse subfamilies

| Expansion | Pouched<br>rat OR<br>family | Pouched rat<br>OR<br>subfamily | Associated<br>mouse<br>subfamily | Number of<br>Pouched rat ORs | Number of<br>mouse ORs |
| --- | --- | --- | --- | --- | --- |
| 1 | 101 | 329 | 150 | 35 | 6 |
| 2 | 101 | 312 | 137 | 12 | 4 |
| 3 | 101 | 321 | 135 | 15 | 1 |
| 4 | 101 | 317 | 136 | 33 | 5 |
| 5 | 5 | 12 | 89 | 10 | 1 |
| 6 | 72 | 219 | 71 | 7 | 2 |

### Orthologous genes

We found that many pouched rat genes were orthologous to mouse, rat, and squirrel OR genes, using the tree in **Fig 2**. 99 of the 1145 putative olfactory receptor genes shared a 1-to-1 orthology with rat, mouse, and squirrel. Using the percent similarity-method for defining families, 10 of the 99 orthologs came from family 38 (ORA equivalent family: OR 51) and 11 from family 101 (ORA equivalent families: OR 1 and OR 7). This further depicts an unequal distribution of orthologs across all of the 87 families defined using this method.

The average similarities of the associated pouched rat gene to these orthologous genes were 92.4% for mouse, 91.9% for rat, and 86.1% for squirrel. However, in other locations we observed potentially paralogous ORs, and there were multiple pouched rat ORs for one rat, mouse, or squirrel OR. For example, Pouchie948 associated with rat olr1356, mouse olfr15, and OR2C1 (squirrel) (**Fig 5a**). We found a total of 33 locations on the gene tree where either the pouched rat, mouse, rat, or squirrel had two paralogous genes. For 1-to-1 orthology of mouse, rat, squirrel, and pouched rat, the most typical pattern (75/99) observed included the mouse and rat being most closely aligned, followed by the pouched rat gene, then the squirrel OR gene (**Fig 5b**).

**Fig 5.**
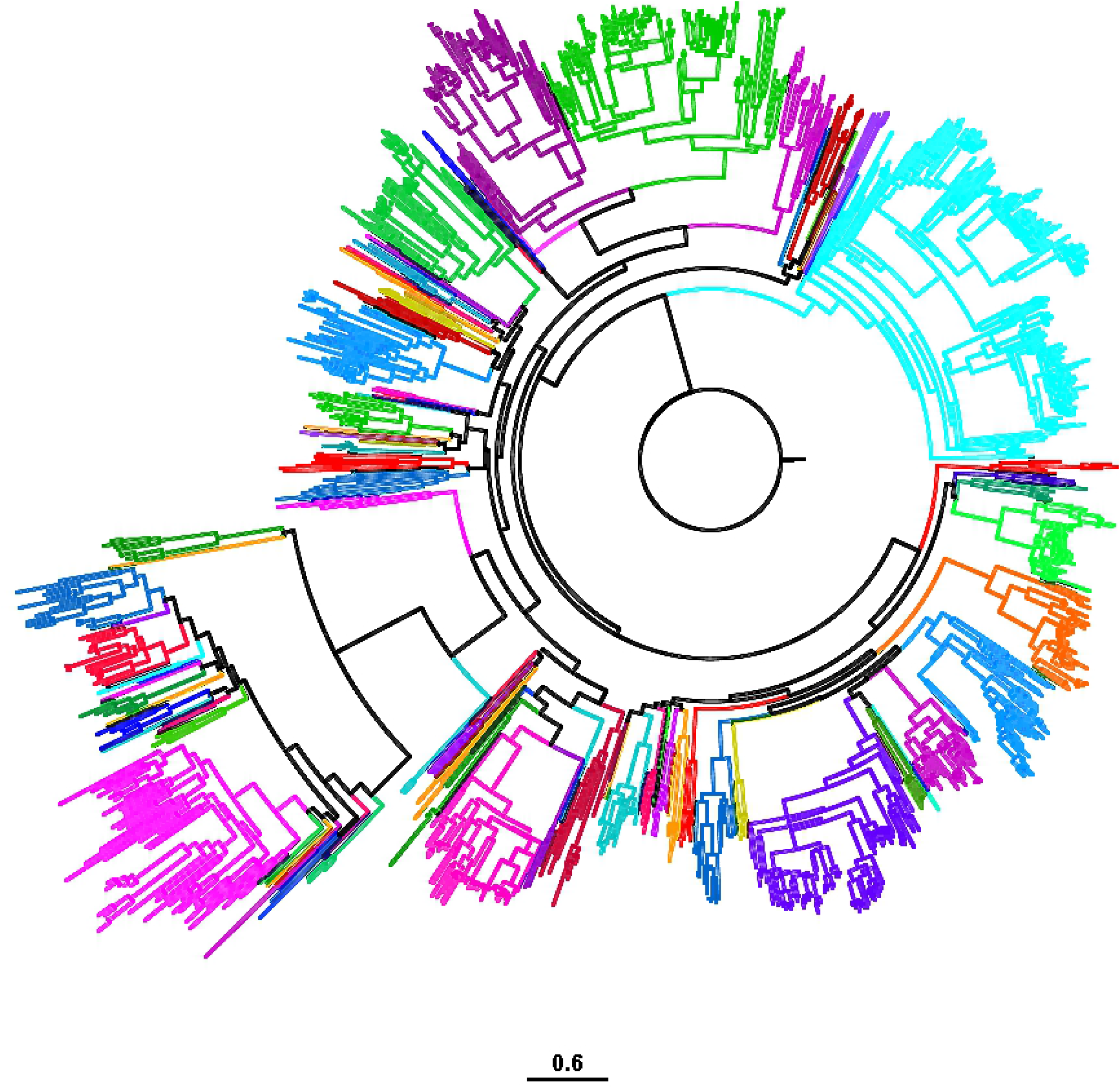
Examples of orthologous olfactory receptor (OR) gene sequence among rodents. Numbers at branch points indicate bootstrap values. A) 1-to-1 orthology between pouched rat, mouse, rat, and squirrel (*Ictidomys tridecemilineatus*) B) Example of pouched rat paralog within orthologous gene grouping

In 33 instances, one or more of the species had a paralogous gene. These paralogous groupings were also very similar, with the pouched rat genes sharing an average of 91.9% sequence similarity to mouse, 91.0% to rat, and 86.2% to squirrel. In 8 cases, there were two paralogous pouched rat OR genes; which had an average pairwise sequence similarity of 97.1%, suggesting a recent duplication of these genes within the pouched rat lineage

166 of 1145 pouched rat genes had 1-to-1 orthology with mouse and rat; all three species are in the Muroidea superfamily. 146 of these orthologies followed the predicted pattern of mouse and rat being closely aligned, followed by the pouched rat gene. Comparatively, there were an additional 44 mouse-pouched rat orthologous genes, 19 rat-pouched rat orthologous genes, and 4 squirrel-pouched rat orthologous genes (**Fig 6**). There were 14 cases of squirrel-mouse-rat orthologous genes. These latter patterns are consistent with differential loss of genes across rodent lineages.

**Fig 6.**
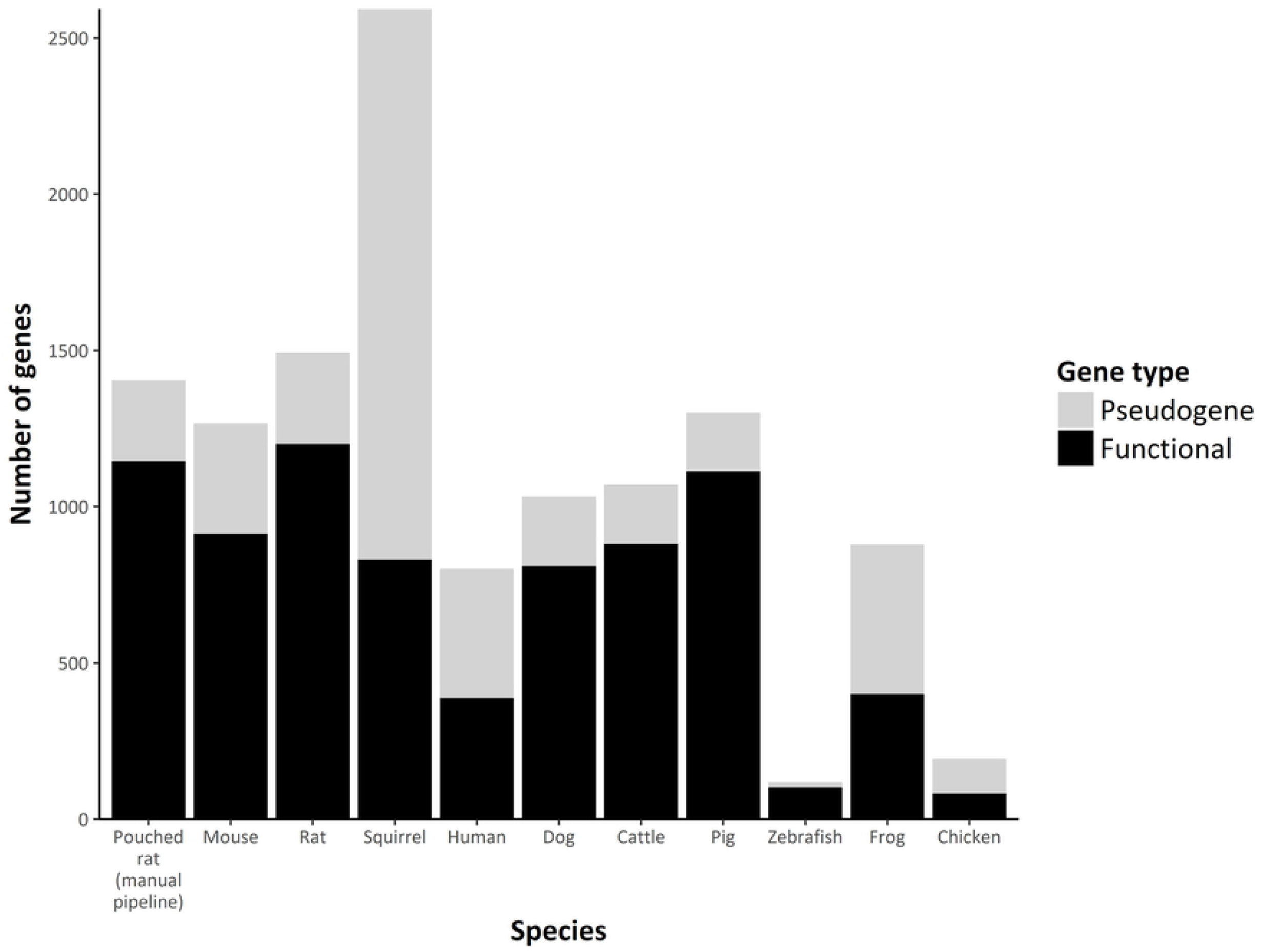
Relative proportions of one-to-one orthology of pouched rat olfactory receptor genes and squirrel, mouse, and rat. Pouched rats have the largest number of orthologous genes shared with mice and rats. The number of orthologous groupings with squirrels are consistently smaller than orthologies with mice or rats, due to phylogenetic placement.

### Clustering of ORs

We examined physical clustering of the ORs based on the locations of the sequences within the gene scaffolds. We found that there were 173 clusters of ORs comprising 644 of the 1145 ORs. All but 50 of these clusters contained ORs that were clustered completely within a family group. The largest clusters contained 12 and 13 sequences (Table 5).

**Table 5.** Distribution of pouched rat olfactory receptors in location clusters

| Number of<br>sequences in<br>cluster | Number of<br>instances |
| --- | --- |
| 2 | 70 |
| 3 | 40 |
| 4 | 22 |
| 5 | 14 |
| 6 | 8 |
| 7 | 5 |
| 8 | 4 |
| 9 | 2 |
| 10 | 3 |
| 12 | 2 |
| 13 | 3 |

We examined where these large clusters were located to determine if the expansions were due to potential duplication events in the genome. 20 of these clusters were located either entirely or partially within these large expansions (Table 6). All of the expansions contained at least one cluster of ORs, supporting the hypothesis that expansions may be partially due to tandem duplications of OR genes.

**Table 6.** Pouched rat olfactory receptor (OR) family expansions and numbers of clustered OR sequences within each expansion

| Expansion | Family | Clusters present (number of sequences represented) |
| --- | --- | --- |
| 1 | 101 | 116 (3 of 4), 154 (4 of 5), 87 (3), 108 (3), 85 (4), 41 (2) |
| 2 | 101 | 72 (2), 168 (3 of 7), 139 (3 of 5) |
| 3 | 101 | 166 (2 of 7), 111 (1 of 2), 147 (3 of 4), 141 (3 of 12), 16 (4 of 9) |
| 4 | 101 | 141 (9 of 12), 151 (1 of 3), 3 (6), 166 (5 of 7), 147 (1 of 4), 16 (5 of 9), 111 (1 of 2), 112 (2) |
| 5 | 5 | 52 (2) |
| 6 | 72 | 92 (1 of 2) |

## Discussion

The olfactory receptor repertoire of the pouched rat is similar to rat and mouse olfactory receptor repertoires but has several lineage-specific expansions. We have shown that these expansions are partially attributable to physically clustered OR genes in the genome, suggesting a role of tandemly arrayed gene duplications in generating additional OR diversity, as has been seen other species (52,55,56). Pouched rats were similar in olfactory repertoire size and pseudogene composition to mice, and rats (Table 2) – which all engage in numerous olfaction-mediated behaviors. Though pouched rats have increased olfactory bulb volume, compared to similar rodents, pouched rats’ olfactory capabilities are mediated by a typical rodent OR repertoire with lineage-specific expansions in several subfamilies.

We had hypothesized that the pouched rats’ olfactory capabilities were supported by anatomical elaboration and an associated genetic expansion of the OR repertoire. In other species, anatomical features including the olfactory bulb size, have been used to effectively predict OR repertoire size (37,60). We observed some subfamily and family-specific expansion, however, the overall number of functional ORs did not differ substantially from mice and rats. The potential advantage of their large olfactory bulbs is unclear – whether these bulbs enable enhanced scent detection for foraging (61), or are an adaptation to being nocturnal (62,63) or territorial (64), or enable better sensitivity to odorants in some way is unknown. Pouched rats may be more ‘specialized’ by having family-specific expansions while maintaining a similar number of functional OR genes compared to mice and rats. Additional investigation of the relationship between olfactory bulb size and OR repertoire in rodents is necessary to determine if other rodents might have enlarged bulbs with typical OR repertoire sizes.

### Expansions in the pouched rat OR gene family

We did not find any evidence that the size of the olfactory subgenome or the number of subfamilies in pouched rat differed substantially from other rodents. The ratio of functional genes to pseudogenes using the manual pipeline search was also similar. It is unclear whether pouched rat pseudogenes may have a retained function (47), or have a loss of function.

As has been reported for other species, lineage-specific expansions tend to occur in tandemly-arrayed clusters (51,52,57). Tandemly-arrayed gene expansions are indicative of a series of gene duplications. We combined overlapping sequences and eliminated duplicates during analysis which could influence the detection of these tandem duplications. Identical sequences within different contigs were eliminated, as we interpreted these as artifacts from assembly. Thus, the number of expansions was conservatively calculated, and future genome assemblies will potentially reveal additional clustering in the genome. Those cases that are shared among mice, rats and pouched rats indicate earlier expansions compared to the tandem arrays specific to pouched rats.

Unfortunately we found no ligands in the literature that were associated with mouse or rat ORs within the same subfamilies of pouched rat OR expansions (14,23,58). Future work on discerning ligands for these ORs might reveal whether pouched rats are primed to specialize in detecting specific types of odorants that are particularly salient and relevant for this species.

### Receptor Conservation among Rodents and within Muroidea

Approximately 9% of the pouched rat olfactory genome shows 1-to-1 orthology with the other rodents in this study – mice, rats and squirrels (**Fig 6**). Pouched rat gene sequences were more similar to mouse and rat sequences than to squirrel sequences, as expected. This agrees with the current phylogenetic hypothesis for rodents: rat and mouse are placed within the Muridae family together, with pouched rat in a different family (Nesomyidae) but within the Muroidea superfamily (**Fig 1**). Squirrels are in a different family (Sciuridae), whereas all four species compared in this study are within Rodentia. Given that pouched rats are more closely related to rats and mice than squirrels are to rats and mice, we should expect the OR phylogeny to reflect this phylogenetic relationship, and it does for these orthologous genes. Further supporting this assertion, an additional 14.5% of the pouched rat olfactory receptor genes have orthologous mouse and rat olfactory receptor genes (**Fig 6**). Pouched rat orthology with only one species was also observed; though this was much more common with either mice or rats compared to the more distant rodent relative – the squirrel.

Interestingly, there were 14 instances where orthology is shared between mouse, rat, and squirrel – potentially pouched rats have lost functional olfactory genes here during their evolutionary history. Four of these potential losses were from family OR 4 (as defined by (11)), although the loss would not substantially affect the number and proportion of functional to non-functional OR genes given the numbers of pseudogenes in the associated pouched rat OR families. Future work investigating pseudogenes may yield additional insight into the orthologous OR genes among rodents. Unfortunately, no ligands have been characterized for these receptors, so we cannot speculate on how this might impact the olfactory capabilities or behavior of the pouched rat.

### Additions to our knowledge about olfaction and OR repertoire

Most rodent species with a described OR repertoire are lab models, and include mice (23) and rats (26), while guinea pigs were included in a comparative study which did not focus on guinea pigs specifically (11). Two other rodent species include some OR information (i.e. an estimate of ORs and pseudogenes based on published genomic information), but it is unclear which search terms were used, and whether the algorithm used would obtain OR families that were not shared by the exemplars (i.e. novel subfamilies). These species included the kangaroo rat (*Dipodomys ordii*) and the thirteen-lined ground squirrel (also included in this study) (11); both of these species are native to North America. Here we have shown that the functional pouched rat OR repertoire is very similar in size to those of other Muroids. The pouched rat OR repertoire is also more conserved within Muroids than within Rodentia, as expected. Our characterization of the orthologous genes within Muroidea suggests that there may be a set of conserved Muroid-specific ORs, although additional species will need to be sequenced and compared to support this hypothesis. The addition of the African giant pouched rat to described rodent OR repertoires also can serve as an outgroup for future analyses given its position in the Nesomyidae family, which has not been previously described.

We noted a pattern in our gene tree of rodent ORs where squirrel and pouched rat ORs tend to branch earlier than the mouse and rat ORs. Given that mice and rats are more closely related to pouched rats than to squirrels, we expected that orthologous genes might follow this pattern. We assume these genes should be orthologous and bind similar odorants with similar structures due to their similar protein sequences, however, it could be that small changes in these genes create ORs that bind different odorants, or change perception in other ways (20,21,59). Identification of these potentially conserved ORs is important for understanding how the OR repertoire might respond to evolutionary pressures over time.

### Conclusions

The description of the African giant pouched rat olfactory receptor repertoire adds to our understanding of how these repertoires vary within Muroidea and among rodents, and how these genes may have been conserved or diverged to serve different behavioral purposes. Our results suggested that there could be a conserved set of ‘Muroidea’ olfactory receptor genes. Whether pouched rats have enlarged olfactory bulbs to enhance their olfactory capabilities in spite of a typical OR repertoire size remains an open, yet interesting question. Further investigation into family-specific expansions within the pouched rat OR repertoire compared to mice and rats might reveal whether these changes have evolved as a way for pouched rats to olfactorily specialize within their habitat.

## Acknowledgments

We wish to thank Andrew Legan for his assistance on this project.

## Supporting Information

**S1 File. Fasta file of all 1145 ORs from the manual pipeline search.**

**S2 File. Fasta file of all ORs from the ORA automated search method.**

**S3 File. Tree file from Pouched rat ORs from manual pipeline search**

**S4 File. Tree file from Rodent ORs, Pouched rat ORs from manual pipeline search**

**S5 File. Tree file from Rodent ORs, Pouched rat ORs from ORA automated search method**

**S6 File. File categorizing family groupings for Pouched rat ORs**

**S1 Table. Distribution of pouched rat olfactory receptor (OR) genes among families using the automated olfactory receptor assigner.**

